# Multicellularity and increasing Reynolds number impact on the evolutionary shift in photo-induced ciliary response in Volvocales

**DOI:** 10.1101/2023.12.22.573154

**Authors:** Noriko Ueki, Ken-ichi Wakabayashi

## Abstract

Volvocales include species with different cell numbers and sizes, reflecting a history of gradual size increase evolution. Unicellular species live in low Reynolds-number (*Re*) environments where viscous forces dominate, whereas multicellular species live in higher *Re* environments with nonnegligible inertial forces. Despite significant changes in the physical environment, during the evolution of multicellularity they maintained photobehaviors (i.e., photoshock and phototactic responses), which allows them to survive under changing light conditions. In this study, we classified photo-induced ciliary responses in Volvocales into four patterns: temporal waveform conversion, no obvious response, pause in ciliary beating, temporal changes in ciliary beating directions. We found that which species exhibit which pattern depends on *Re* associated with the individual size of each species rather than phylogenetic relationships. These results suggest that species with increased cell numbers acquired their responses adapted to higher *Re* fluid environments.

**Significance Statement:** Volvocales green algae include species with various cell numbers and are excellent organisms for studying the evolution of multicellularity. They exhibit photobehaviors by changing the pattern of ciliary beating, which could be categorized into four patterns. We found that the difference in patterns among the organisms is due to the Reynolds number, the ratio of viscous and inertia forces, rather than their phylogenetic relationships. This study indicates that the fluid environment was an important factor in natural selection for behavioral changes in microalgae during evolution. The results link evolution and physics while contributing to the design of micromachines.

## Introduction

In the history of evolution, multicellularity occurred multiple times independently; multicellularity is thought to result in survival advantages under selective pressure (1). Multicellularity provides various benefits: avoiding predation due to larger size, effective predation/escape/movement to optimal environments due to faster moving speed, division of labor by cell differentiation, etc.

Volvocales green algae are a group of swimming phytoplankton that include species with a wide range of cell numbers, making them an excellent model for studying the evolution of multicellularity in existing organisms (Fig. 1A). The multicellular species in Volvocales have either a flat or spherical shape. Flat species contain relatively few cells, like *Tetrabaena* and *Gonium* (Fig. 1A, 1C). Among spherical species, *Pandorina* has a morphology of densely packed cells, and those with more cells have a single layer of cells on the surface of the spheroid, like *Eudorina*, *Pleodorina*, and *Volvox* (Fig. 1A, 1C). Their evolution involved increasing complexity, such as in morphogenesis and sexual reproduction (2). *Volvox*, which has the highest complexity, has the following survival advantage traits: they avoid predation by their large size (3, 4), swim fast (5), move quickly to optimum light environments (6), and differentiate between somatic and reproductive cells, with the reproductive cells being located and protected inside the spherical somatic cell sheet (7); Fig. 1A).

**Figure 1.**
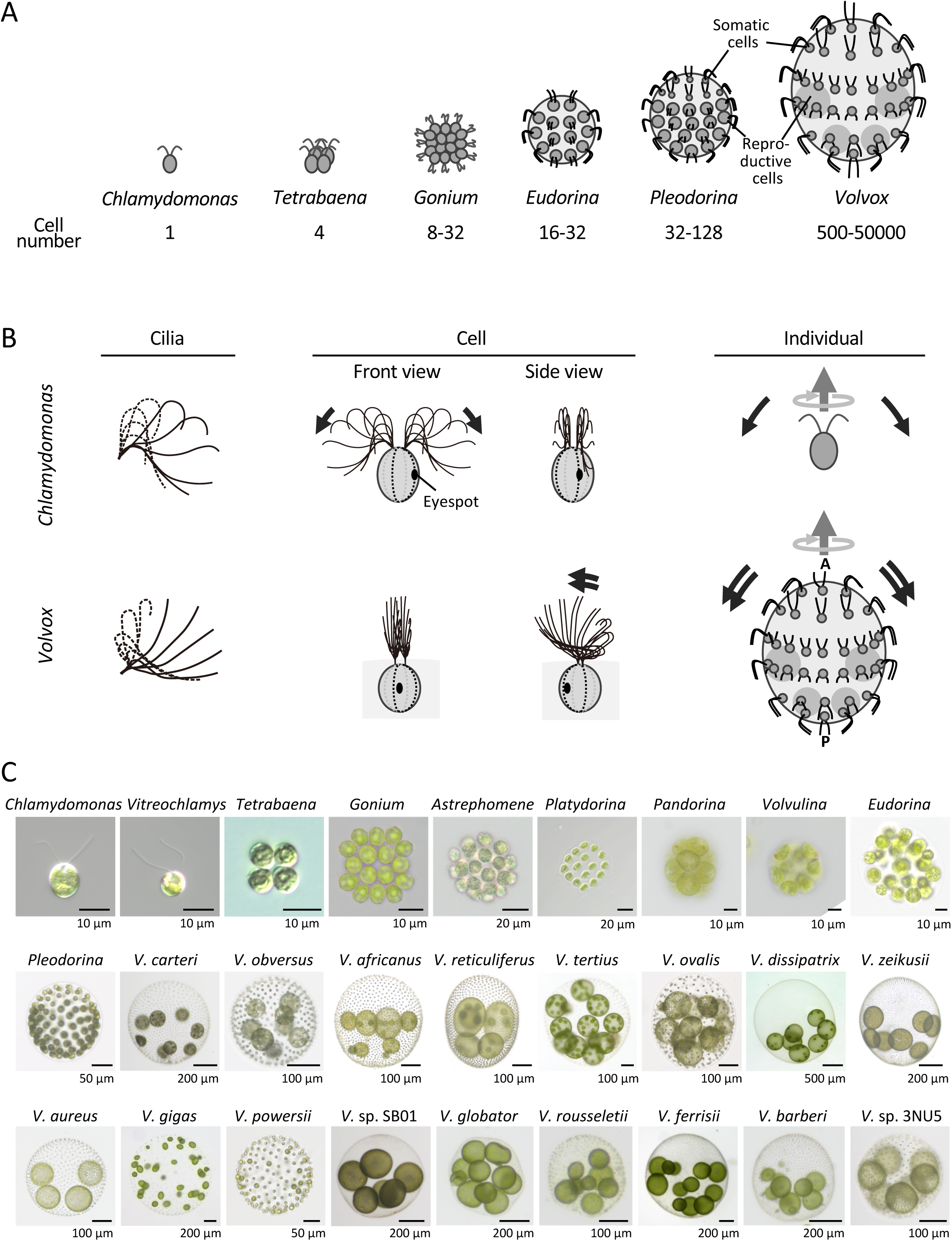
Configuration of cell and cilia in Volvocales. (A) Schematic illustration of representative genera in Volvocales. (B) Ciliary and individual movement in *Chlamydomonas* and *Volvox*. Left: one stroke of an asymmetric waveform of cilia. Effective stroke (solid lines) and recovery stroke (broken lines) are shown. Modified by (33) and (6). Middle: Relationship between a cell and the direction of ciliary beating. Right: direction of the individual’s forward swimming and rotation. Black arrows: direction of ciliary beating in the asymmetric waveform. Dark gray arrows: individual’s swimming direction. Light gray arrows: direction of rotation. A, anterior pole; P, posterior pole. (C) Species/strains in Volvocales used in this study. Note the difference in scale bars. For species other than *Volvox*, only generic names are shown. For *Volvox*, abbreviated generic names and specific names are shown. See Table 1 for details.

**Table 1.**
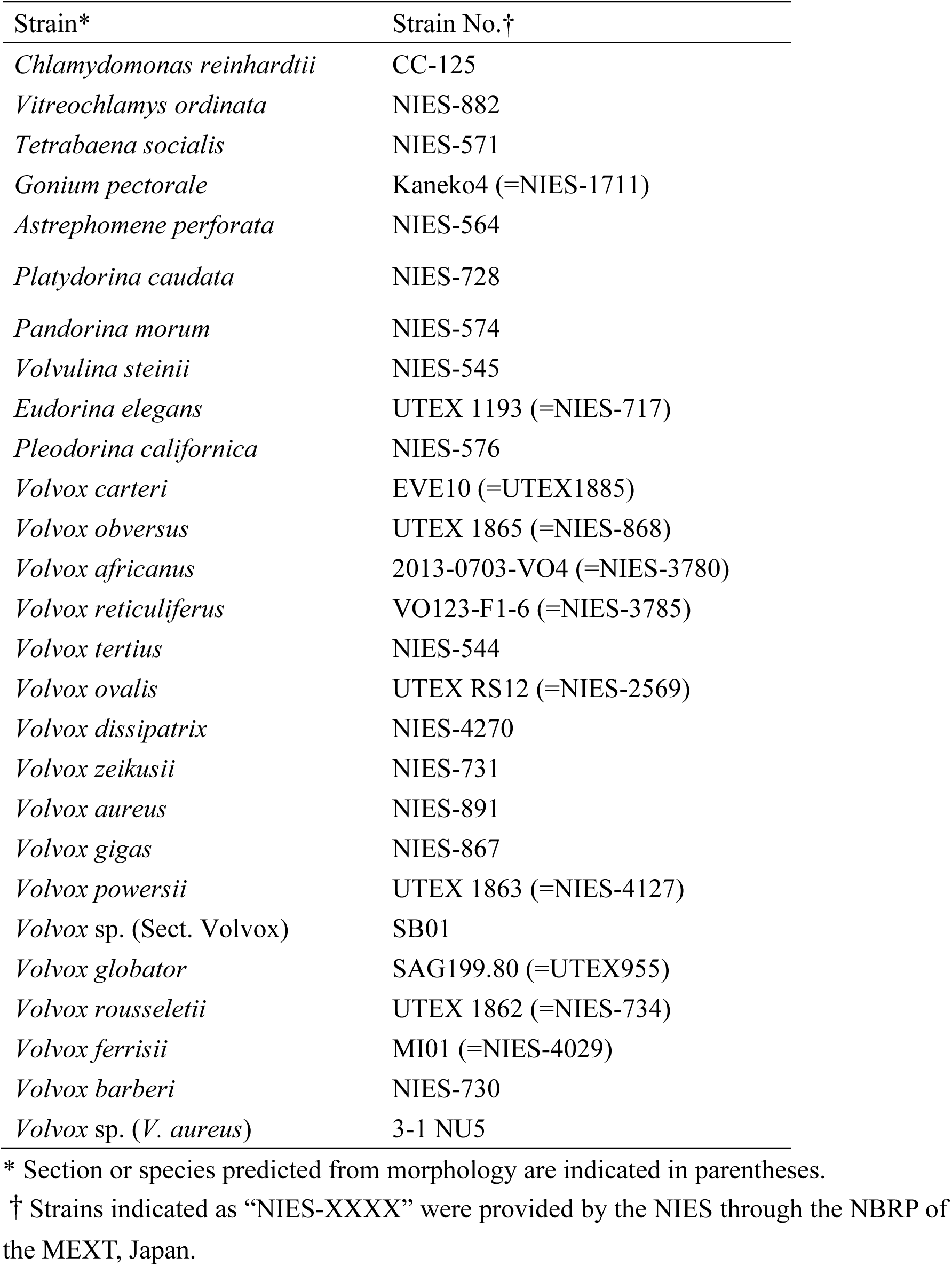
Volvocales strains used in this study.

With increasing size of aquatic organisms, the physical properties of the surrounding fluid change. For organisms as small as 10 µm, such as *Chlamydomonas*, the environment is dominated by viscous forces. This would be like humans swimming in honey; thus, those small organisms must constantly generate propulsive forces to move. Cilia (or flagella) are considered adequate in such environments (8). For smaller organisms ∼1 µm in diameter, such as *Escherichia coli,* the effect of diffusion by Brownian motion cannot be ignored. For larger ones, >1 mm, such as *Volvox* or *Daphnia*, the effect of inertia cannot be ignored (9, 10). Accordingly, Volvocales range from sizes where viscous forces are dominant to those with nonnegligible inertial forces. The ratio of inertia force to viscosity force is defined by the Reynolds number (*Re*): for *Re* > 1, inertia overcomes viscosity.

Volvocales species commonly possess one eyespot and two cilia on each cell, and beat the cilia toward the posterior of the individual with asymmetric waveforms consisting of effective and recovery strokes (Fig. 1B) (11). Because the cilia deform differently during the two strokes, propulsion is efficiently generated even in environments with low *Re*, i.e., where viscosity is dominant (12). When the eyespot senses a light stimulus, Ca^2+^ flows into the cilia and regulates ciliary movement (13). Interestingly, several reports suggest that the regulation pattern in ciliary movement varies among species. In *Chlamydomonas reinhardtii*, the ciliary waveform converts from asymmetric to symmetric in response to light, causing temporal backward swimming (14–17). In *Tetrabaena socialis*, which has a flat shape with four cells, cilia do not change their motility even after light stimulation, and no photobehavior is observed (18). Several *Volvox* species temporarily reduce the ciliary beating frequency or stop ciliary movement in response to light (19). Other *Volvox* species in section Volvox, a group containing relatively more cells characterized by thick cytoplasmic bridges connecting cells, cilia temporarily change the direction of beating in response to light while remaining into an asymmetric waveform [(6) for *V. ferrisii,* previously reported as *V. rousseletii*; (19) for *V. barberi*].

Thus, Volvocales species commonly beat their cilia in an asymmetric waveform toward the backward of the individual during normal forward swimming; conversely, the responses of ciliary movement after light differ among species. Here, the four patterns above are classified according to the number of cells and size of the individual as “1: temporal waveform conversion,” “2: no obvious response,” “3: pause in ciliary beating,” and “4: temporal changes in ciliary beating directions”. In this study, we categorized the pattern of photo-induced ciliary response for 27 strains (Fig. 1C) and found that the patterns matched better with the *Re* of each species than with phylogenetic relationships.

## Results

### Photo-induced ciliary responses could be categorized into four patterns in Volvocales

To understand the diversity of ciliary responses to light in Volvocales, we observed ciliary movements after exposure to a white flashlight in 27 strains. The response mode was categorized into the following four patterns (summarized in Fig. 2E). Hereinafter, for readability, non-abbreviated generic names are shown for species other than *Volvox*.

**Figure 2.**
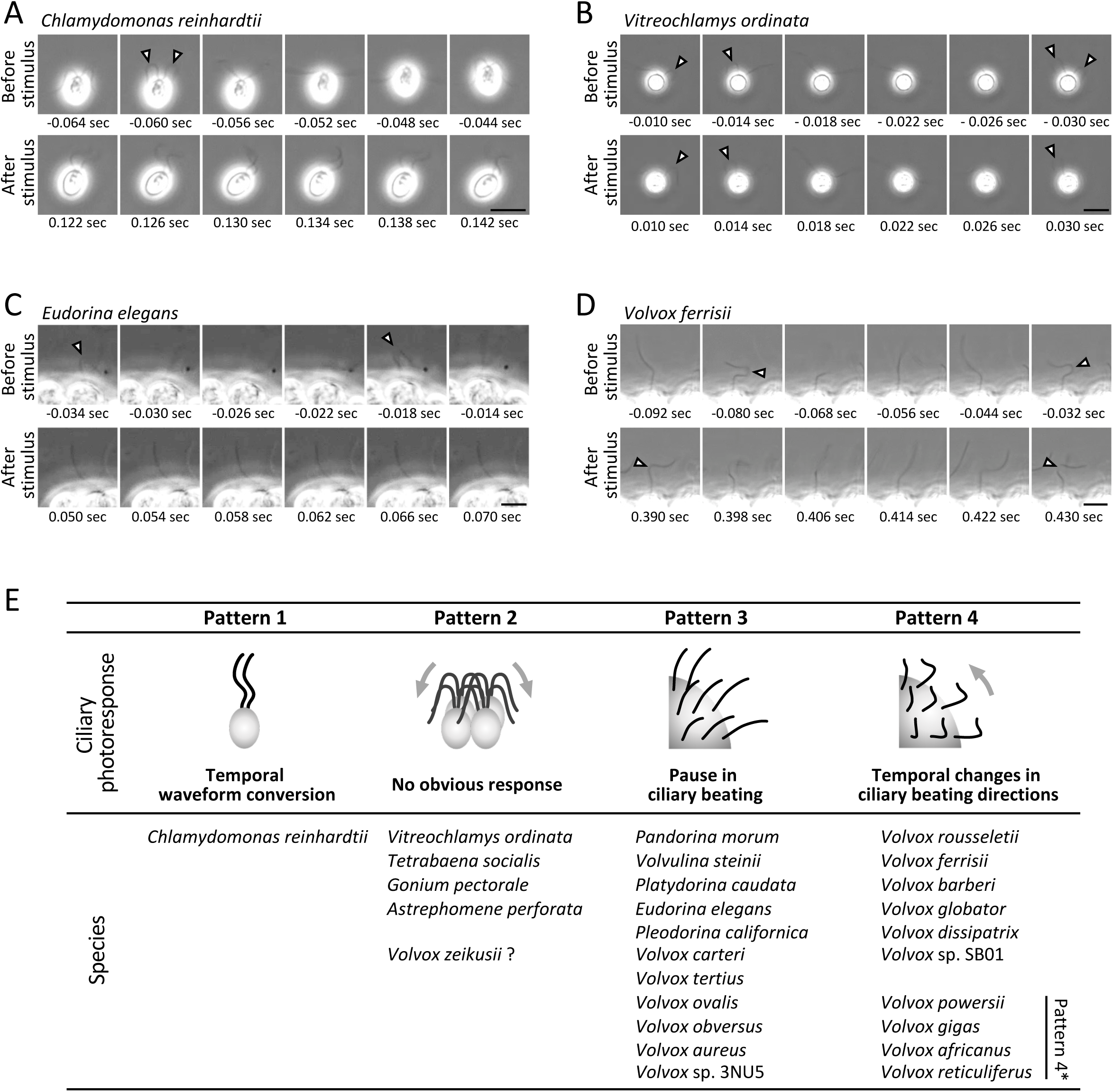
Four patterns of photo-induced ciliary response. (A–D) Upper rows: sequential images at equal intervals covering >1 stroke of the asymmetrical waveform before flashlight application. Bottom rows: sequential images after flashlight application in the same time interval as the upper rows. Time points from flashlight application are shown. White arrowheads indicate bends of asymmetric waveforms. (A) Temporal waveform conversion (Pattern 1) in *Chlamydomonas reinhardtii*. (B) No obvious response (Pattern 2) in *Vitreochlamys ordinata*. (C) A pause in ciliary beating (Pattern 3) in *Eudorina elegans*. (D) Temporal changes in ciliary beating directions (Pattern 4) in *Volvox ferrisii*. Scale bars: 10 μm. (E) Four patterns of photo-induced ciliary response. Volvocales species exhibiting each pattern are shown. Asterisk indicates species exhibiting Patterns 3 and 4.

#### Pattern 1: Temporal waveform conversion

In the unicellular species *Chlamydomonas*, cilia transiently converted their waveform from asymmetric to symmetric immediately after light stimulation (Fig. 2A and E and Movie S1). Only *Chlamydomonas* showed this pattern among the species examined.

#### Pattern 2: No obvious response

No obvious light response has been reported for *Tetrabaena* (18). Here, three closely related species, unicellular *Vitreochlamys*, 8–16 cells *Gonium,* and 16–32 cells *Astrephomene*, did not obviously respond to the light stimulus (Fig. 2B and E and Movie S2). In *Vitreochlamys,* spontaneous conversion from asymmetric to symmetric waveforms was frequently observed independent of light stimulation, as reported for *Tetrabaena* (18). In addition, *V. zeikusii*, a lineage far apart from the small species above, did not respond to the light stimulus.

#### Pattern 3: Pause in ciliary beating

Eleven strains of Volvocales temporarily ceased ciliary movement in response to light stimulation (Fig. 2C and E and Movie S3). In addition, cells closer to the anterior pole of the individuals were more responsive (Movie S3). *Pandorina* and *Volvulina* showed relatively short cessations of around 0.05 s, while larger species, such as *Pleodorina,* showed rather long arrest durations (several seconds).

#### Pattern 4: Temporal changes in ciliary beating directions

Relatively larger species exhibited temporal ciliary reversal or change in the beating direction of an asymmetric waveform, all of which are *Volvox* (Group 4; Fig.2D and E and Movie S4). Similar to the organisms that exhibit Pattern 3, the cells closer to the anterior pole were more responsive. All species of the section Volvox used in this study (*V. rousseletii*, *V. ferrisii*, *V. barberi*, and *V. globator*; Fig. 3, and *Volvox* sp. SB01) and *V. dissipatrix* which is sampled in Thailand with a maximum diameter greater than 2 mm (20) exhibited only this pattern. The response typically lasted for one to several seconds. In the section Volvox, the asymmetric waveforms in the changed direction during the response were narrower in amplitude and more three-dimensional than the usual asymmetric waveforms, as previously reported (6). In contrast, outside of the section Volvox, *V. dissipatrix*, the shape and the amplitude of asymmetric waveform during the response were almost identical to those before light irradiation. The time lag between light application and the onset of the ciliary response was ∼0.2 s in the section Volvox. *V. dissipatrix* tended to exhibit a longer time lag (∼1.5 s).

**Figure 3.**
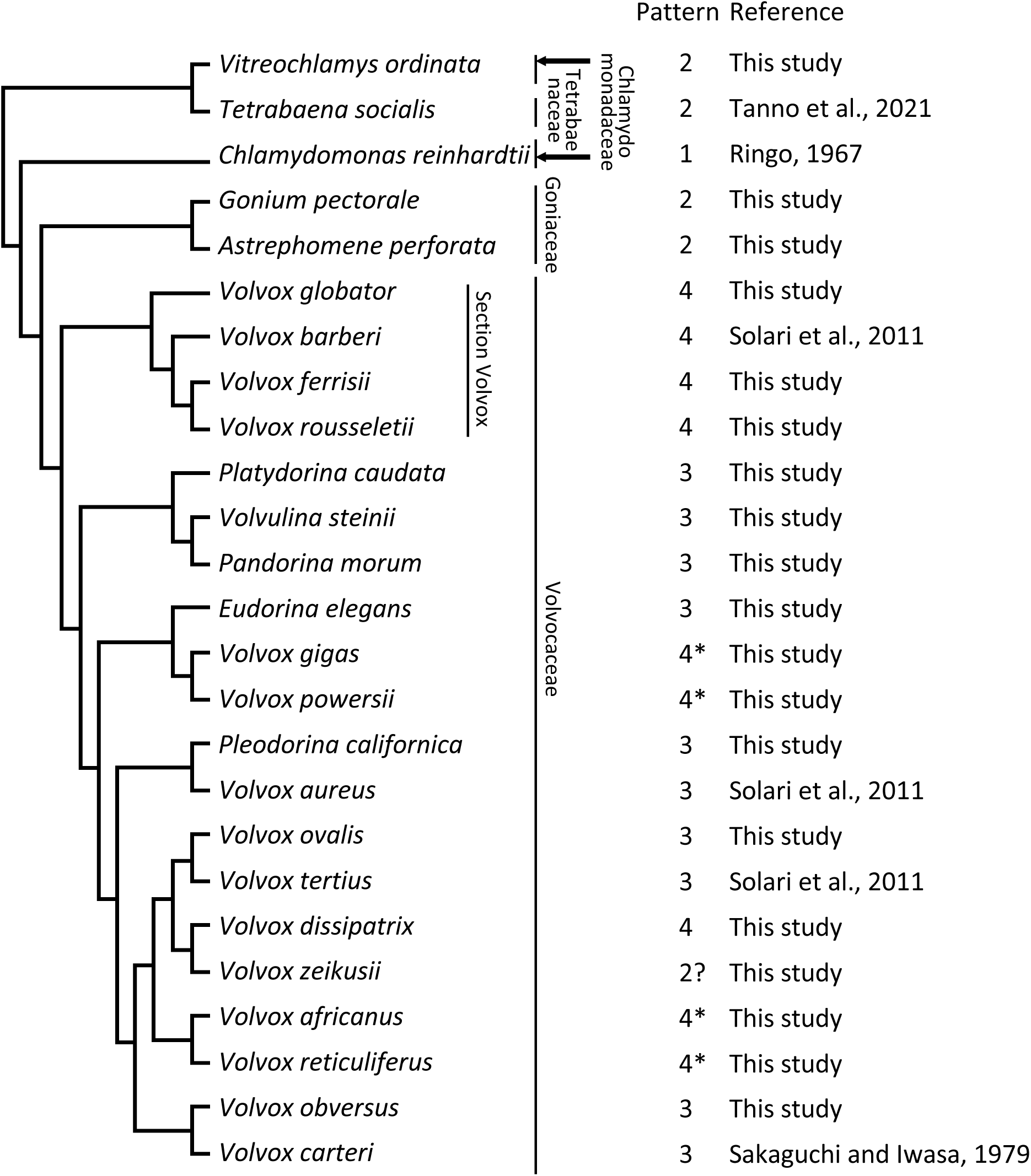
Mapping of the four patterns of photo-induced ciliary response onto the phylogenic tree of species used in this study. Phylogenetic relationships are based on (34). Asterisks indicate species exhibiting Patterns 3 and 4. Bar length indicates the order of branching, not its timing.

In addition, several other species exhibited both Pattern 3 and Pattern 4, and we termed this pattern Pattern 4* (Fig. 2E). In *V. powersii*, *V. reticuliferus*, and *V. africanus*, both patterns were observed in different areas of an individual. In *V. gigas*, the ciliary movement stopped immediately after light stimulation, followed by beating with an asymmetric waveform that changed direction by about 90° then returning to the original beating direction.

### Photo-induced ciliary responses are not completely branch-dependent

The phylogenetic relationship of species used in this study known thus far and their patterns of photo-induced ciliary response are shown in Fig. 3. The family Volvocaceae is distinguished from the other three families by its spherical shape and inversion during development. All strains in Volvocaceae exhibited Pattern 3 or 4, or both. All other phylogenetic groups showed Pattern 1 or 2. Pattern 1 was observed only in the unicellular *Chlamydomonas.* Pattern 2 was observed in the unicellular *Viotreochlamys* and multicellular species in Tetrabaenaceae and Goniaceae. All species in section Volvox and *V. dissipatrix*, phylogenetically distant from section Volvox, showed only Pattern 4. *V. gigas* and *V. powersii*, and *V. africanus* and *V. reticuliferus*, which are closely related to each other, showed both Patterns 3 and 4.

### Relationship between cell number and size and patterns of photo-induced ciliary responses

The above data suggest that the patterns of photo-induced ciliary response are not solely dependent on lineage. We surmised that the number of cells in an individual and the size of the individual may be factors determining the pattern. To examine this possibility, the number of cells and anterior-posterior length of the largest single individual in each strain’s culture were measured and plotted (Fig. 4). The four patterns of photo-induced ciliary response are distinguished by symbols. Pattern 1 is exhibited by only unicellular *Chlamydomonas*, Pattern 2 was found in individuals with 1–64 cells (and 10,362 cells for *V. zeikusii*), Pattern 3 in those with 16–3,037 cells, Pattern 4* in individuals with 1,057– 2,813 cells, and Pattern 4 in those with 3,788–23,422 cells. With respect to anterior-posterior length, Pattern 1 was associated with a length of 11 µm, Pattern 2 to one between 30–130 µm (1000 µm for *V. zeikusii*), Pattern 3 with lengths of 42–1,780 µm, and Pattern 4 for sizes of 500–2,230 µm. Despite overlaps, we identified a rough transition from Pattern 1 to 4 according to cell number and size.

**Figure 4.**
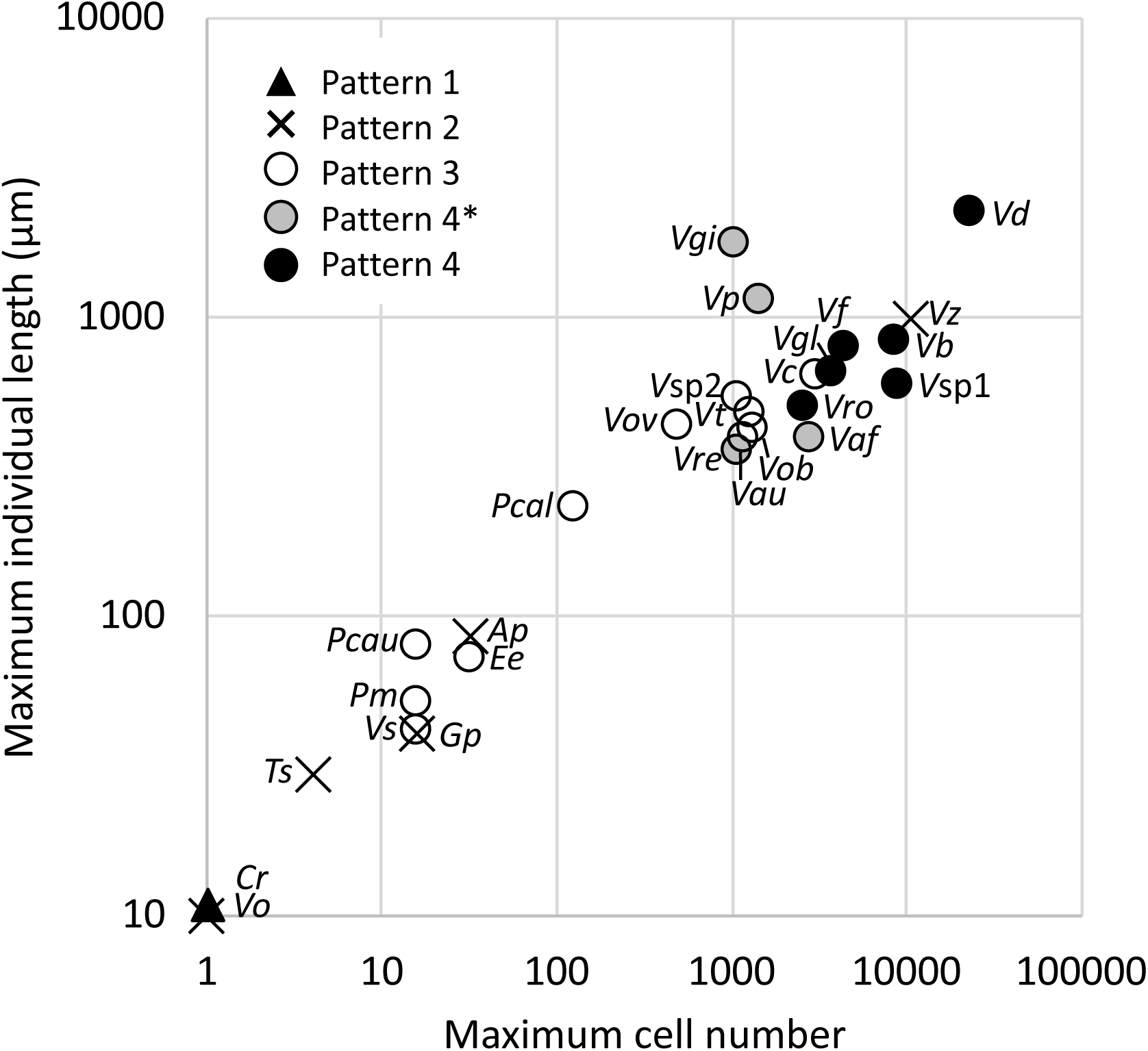
Relationships among maximum cell number of individuals, maximum length along the anterior-posterior axis of an individual, and the patterns of photo-induced ciliary response in each species. A black triangle indicates Pattern 1, crosses indicate Pattern 2, white circles indicate Pattern 3, and gray and black circles indicate Pattern 4. Gray circles indicate species that exhibit Patterns 3 and 4. Data on *Tetrabaena* is derived from (18). Species are denoted by the first letter of genus name and the first 1-3 letters of specific name. *V*sp1: *Volvox* sp. SB01; *V*sp2: *Volvox* sp. 3-1 NU5. See also Table 1.

### Relationship between Reynolds number and patterns of photo-induced ciliary response

How and why cell number and size relate to the patterns of photo-induced ciliary response were examined in terms of the surrounding physical environment, i.e., fluid. Fluids are characterized by the ratio of viscous and inertial forces, the Reynolds number (*Re*), which can be calculated using the diameter of the organism in swimming direction and swimming velocity. In other words, the fluid environment is determined by the individual size and the driving force which depends on the number of cells with two cilia each.

We calculated the *Re* for these organisms, and found that this value can be roughly associated with their patterns of photo-induced ciliary response: around 10^−3^ in Group 1, mainly 10^−3^ to 10^−4^ in Group 2, mainly 10^−2^ to 10^−1^ in Group 3, and mainly 10^−2^ to 10^0^ in Group 4 (Fig. 5).

**Figure 5.**
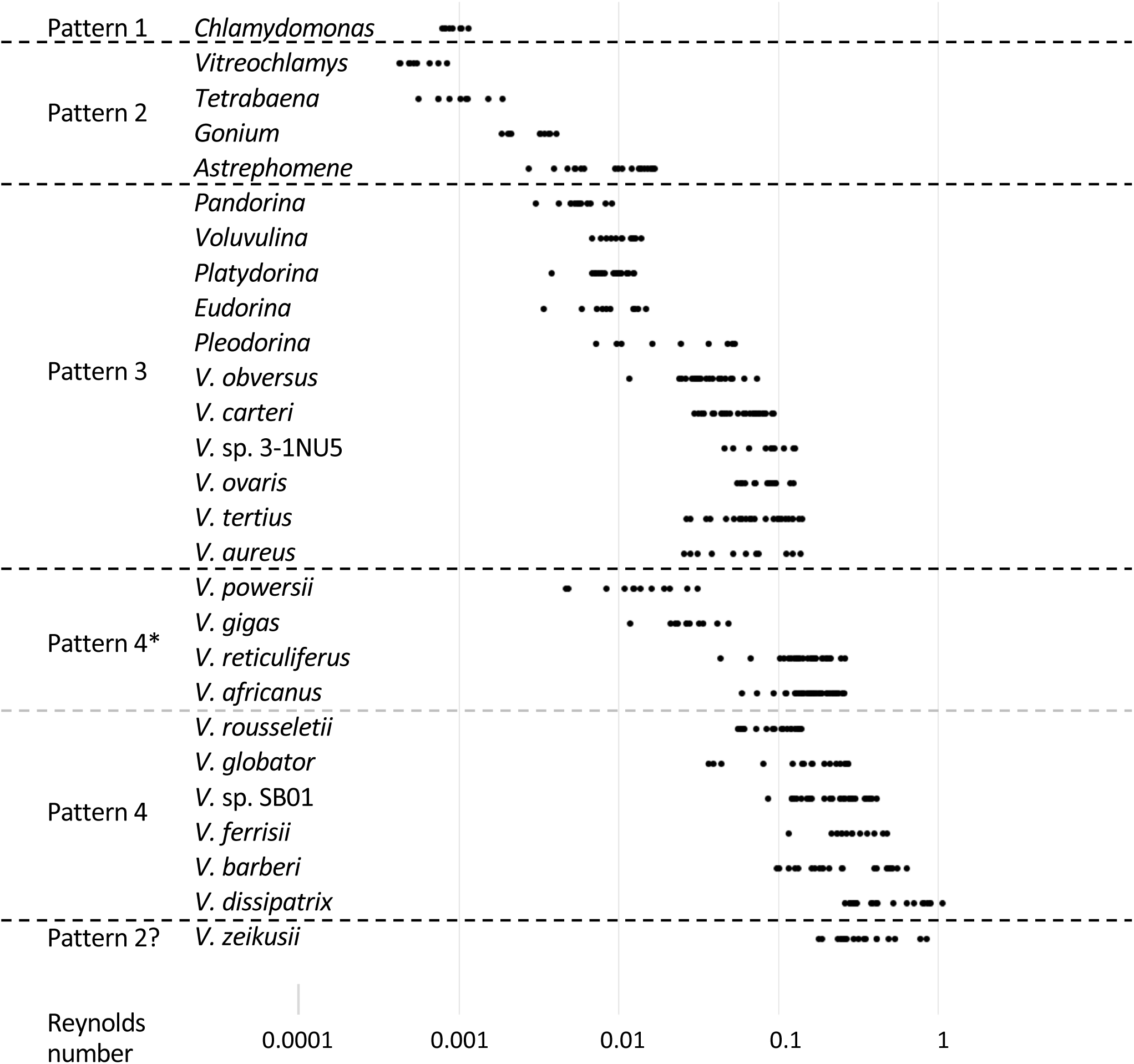
Reynolds numbers of 27 strains in Volvocales for each pattern. Each dot derives from an individual.

## Discussion

In this study, we observed four distinct patterns of photo-induced ciliary response among 27 Volvocales strains. They were roughly dependent on cell size, cell number, and *Re*.

### A possible unique strategy of early-branched species in Volvocales that exhibited “no obvious reaction”

Pattern 2, no obvious response, was shared by *Vitreochlamys*, *Tetrabaena*, *Gonium*, and *Astrephomene* (Fig. 2E and Fig. 3). *Tetrabaena* has been reported to have chloroplasts with high photoprotection ability, and its eyespot does not function as a photoreceptor apparatus despite an intact eyespot structure like that of *Chlamydomonas* (18) (21). This is considered as a strategy to avoid high-light stress, in contrast to that in *Chlamydomonas*, which quickly escapes from light. The same strategy could be shared by species exhibiting Pattern 2.

In response to a flashlight, although both are unicellular, *Chlamydomonas* exhibited Pattern 1, waveform conversion, as previously reported (22), while *Vitreochlamys* did not respond obviously to a flashlight. Considering that laboratory strains of *Chlamydomonas* were originally isolated from soil (23) and *Vitreochlamys* from a lake (24), *V. ordinata* may have different survival strategy from that of *Chlamydomonas*: i.e., gaining photoprotection ability like *Tetrabaena*, because of the high-light environment of the lake. This idea should be confirmed by analyses of photosynthetic parameters in *Vitreochlamys*.

Among the species showing Pattern 2, *Tetrabaena* has been reported to exhibit no phototaxis (18), and *Gonium* to do so (25). In these species, it is likely that the photoshock response to a flashlight and phototaxis to a directional continuous light are caused by different mechanisms, as for *Chlamydomonas*, for which mutants exhibiting only photoshock response or phototaxis have been isolated (26, 27).

### Cilia motility changed significantly with the evolution to Volvocaceae

Volvocaceae’s Patterns 3 and 4, pause and direction change, are distinctly different from those shown by early-branched species in Volvocales. The following changes might have occurred during the evolution to Volvocaceae. (1) The eyespot’s function as a photoreceptor, which appears to be lost in Pattern 2 species, became functional again. (2) Symmetrical waveforms were observed in response to light in Pattern 1 species and independent of light in Pattern 2 species [*Tetrabaena* (18) and *Vitreochlamys* (this study)]. Conversely, symmetrical waveforms in species with Patterns 3 and 4 have not been observed, suggesting that these organisms have lost the trait of cilia beating in symmetrical waveforms. (3) Instead, new Patterns 3 and/or 4 are acquired: high intraciliary Ca^2+^ concentration induces a transient pause or direction change instead of the symmetric waveform. (4) The photoprotection ability at the same level as Pattern-2 species can be maintained. (5) The multicellular spherical shape probably inevitably led to the patterns of photo-induced ciliary response in both photoshock response and phototaxis. The use of ciliary response for an individual’s photobehaviors has been demonstrated in *V. ferrisii* (6).

### Direction-change pattern is related to an increase in Reynolds number associated with multicellularity

Volvocaceae species exhibited Pattern 3 and/or 4, which are not completely branch-dependent (Fig. 3) suggesting that Pattern 4 may have been independently acquired multiple times. The transition from Pattern 3 to Pattern 4 seems to occur when the cell number exceeds several thousand, and the anterior-posterior length of individuals exceeds about 600 µm (Fig. 4). Pattern 4 species, which have higher cell number and size also tended to have higher *Re* (Fig. 5). All species that exhibit Pattern 4 showed a maximum *Re* > 0.1, reaching 1 in *V. dissipatrix*. At those *Re*, it is likely that the individual’s movement could no longer be controlled by the ciliary pause of Pattern 3, due to increased inertia. Instead, it appears that the movement is controlled by the direction change of Pattern 4, creating a water flow in the opposite direction.

The structure and mechanism in cilia that enable the alteration in the ciliary beating patterns remain unclear. Studies using detergent-extracted cell models or isolated cilia showed Pattern 1 in *Chlamydomonas* and Pattern 4 in *V. ferrisii* in demembranated ciliary axonemes by the addition of ATP and Ca^2+^ *in vitro* (Bessen et al., 1980; Ueki and Wakabayashi, 2018). This suggests the molecular mechanism that enables photo-induced ciliary responses exists in a detergent-insoluble fraction, such as ciliary axonemes. Microstructure in the axonemes, which would include proteins that regulate the activity of the motor protein dynein, may differ in the cilia of the four groups of organisms.

### Prospects for the relationship between the ciliary response pattern and *Re* in other organisms

A limitation of this study lies in the inability to distinguish between increases in the cell number and those in *Re*. Since all species with a large number of cells in Volvocales are spherical, and since the propulsive force per cell is considered almost the same, an increase in the number of cells almost directly implies an increase in *Re*. Our results clearly indicate that the pattern of ciliary response is phylogeny-independent, but it is still unclear whether the pattern change is due to the increase in the cell number or those in *Re*.

This issue would be resolved by analyzing the ciliary response of phylogenetically distant organisms from Volvocales. Changes in ciliary beating patterns in response to stimuli are observed in organisms with ciliary movements in general (28). For example, a ciliate *Paramecium* exhibits Pattern 4 upon mechanical stimuli (29). Interestingly, *Re* of *Paramecium* is estimated to be ∼0.1 (10), which is the value of the transition from Pattern 3 to Pattern 4 (Fig. 5). Comparison with the ciliary responses occurring in organisms other than Volvocales, e.g., ciliates and mollusk larvae, will provide a detailed picture of the relationship between ciliary response and *Re*.

In conclusion, we found that diverse regulatory mechanisms of cilia in closely related organisms in Volvocales and that this diversity could be explained by *Re* that increased during the evolution of multicellularity. This study suggests the importance of fluid dynamics as a selection pressure and also provides useful information for the design of propulsion systems in biomimetic micromachines.

## Materials and Methods

### Strains and cultural conditions

The Volvocales strains used in this study are listed in Table I. *Chlamydomonas reinhardtii* strain CC-125 was grown in ∼130 mL of tris-acetate-phosphate medium (30) in a 200-mL flask with aeration at 23°C under continuous white, fluorescent light at ∼120 µmol photons m^−2^ s^−1^. Other strains were grown in *Volvox* thiamin acetate (VTAC) medium (31, 32), except for *Tetrabaena socialis* in artificial freshwater-6 (AF-6) medium (32), statically in test tubes with a volume of 10 mL or in a 200-mL flask with a volume of ∼130 mL at 24°C–28°C on a 16 h/8 h light/dark cycle under white fluorescent light at ∼120 µmol of photons m^−2^ s^−1^.

### Observation of photo-induced ciliary response

Photo-induced ciliary responses were observed under a phase-contrast microscope (Axioscope 2, ZEISS or Axioscope 5, ZEISS) under red illumination (>665 nm). Then, algae were illuminated by a white flashlight (Speedlite 300EZ, Canon). The ciliary beating was video recorded at 500 frames per second using a high-speed CCD camera (HAS-U2, DITECT Corporation).

### Measurement of individual size and counting of cell number of individuals

Individuals of *Volvox* species were placed on a glass slide without a cover slip on top to avoid crushing or changing its size, and were photographed focusing on the periphery and surface of the spheroid using a stereomicroscope (Stemi508, ZEISS) equipped with a digital camera (WRAYCAM-NOA630B, Wraymer). The length in the anterior-posterior direction of the largest individual in the largest developmental stage was measured focusing on the periphery and the number of somatic cells per area was counted focusing on the spherical surface; then, the total cell count was calculated. Species other than *Volvox* were photographed using a digital camera (Zeiss Axiocam 208 color) equipped with an optical microscope (Axioscope 5, ZEISS); the length in the anterior-posterior direction was measured and the total cell number counted in the largest individual.

### Measurement of diameter and swimming speed and Reynolds-number calculation

About 8 mL of culture was placed into a 3.5 cm plastic petri dish (Iwaki) placed on the stage of a stereomicroscope (Stemi508, ZEISS) and videorecorded for around 1 min using an Axiocam 208 color camera (ZEISS). The distances that individuals traveled for 3–10 s were measured using the plugin “Manual Tracking” of FIJI (ImageJ) software version 2.1.0. Velocities were determined based on the time and distance traveled. The diameters of the individuals were measured in the direction perpendicular to the anterior-posterior axis using FIJI (ImageJ) software version 2.1.0. The *Re* for each individual was calculated according to the following formula: diameter (µm) × velocity (µm/s) / kinematic viscosity coefficient for water 10^6^ (µm^2^/s).

## Acknowledgments

We are grateful to Dr. Hisayoshi Nozaki for providing Volvocales strains and for critical suggestions about the selection and culture of strains. We thank the NIES collection for advice regarding culture. This work was supported by Japan Society for the Promotion of Science KAKENHI Grants to NU (19K23758, 21K06295) and KW (22H02642,

22H05674, 22H01440, 23K18136), by Ohsumi Frontier Science Foundation to KW, and by Dynamic Alliance for Open Innovation Bridging Human, Environment and Materials to NU and KW.

