## Supplemental text for "Multicellularity and increasing Reynolds number impact on the evolutionary shift in photo-induced ciliary response in Volvocales"

**Supporting Information**

**Movie S1.**

Movie for Fig. 2A, showing a *Chlamydomonas reinhardtii* cell photostimulated at 2.7 s from the start of the video. The movie is run at ×1/17 speed.

**Movie S2.**

Movie for Fig. 2B, showing a *Vitreochlamys ordinata* cell photostimulated at 7.1 s from the start of the video. The movie is run at ×1/17 speed.

**Movie S3.**

Movie for Fig. 2C, showing a *Eudorina elegans* spheroid photostimulated at 3.1 s from the start of the video. The arrows indicate the cilia shown in Fig. 2C. The movie is run at ×1/17 speed.

**Movie S4.**

Movie for Fig. 2D, showing a *Volvox ferrisii* partial spheroid near the anterior pole photostimulated at 4.9 s from the start of the video. The asterisk indicates the cell shown in Fig. 2D. The movie is run at ×1/17 speed.
